# Luteinizing hormone stimulates ingression of mural granulosa cells within the mouse preovulatory follicle

**DOI:** 10.1101/2023.04.21.537855

**Authors:** Corie M. Owen, Laurinda A. Jaffe

**Author notes:** Correspondence: Department of Cell Biology, University of Connecticut Health Center, 263 Farmington Ave., Farmington, CT 06030 USA.,.

## Abstract

Luteinizing hormone (LH) induces ovulation by acting on its receptors in the mural granulosa cells that surround a mammalian oocyte in an ovarian follicle. However, much remains unknown about how activation of the LH receptor modifies the structure of the follicle such that the oocyte is released and the follicle remnants are transformed into the corpus luteum. The present study shows that the preovulatory surge of LH stimulates LH receptor-expressing granulosa cells, initially located almost entirely in the outer layers of the mural granulosa, to rapidly extend inwards, intercalating between other cells. The cellular ingression begins within 30 minutes of the peak of the LH surge, and the proportion of LH receptor-expressing cell bodies in the inner half of the mural granulosa layer increases until the time of ovulation, which occurs at about 10 hours after the LH peak. During this time, many of the initially flask-shaped cells appear to detach from the basal lamina, acquiring a rounder shape with multiple filipodia. Starting at about 4 hours after the LH peak, the mural granulosa layer at the apical surface of the follicle where ovulation will occur begins to thin, and the basolateral surface develops invaginations and constrictions. Our findings raise the question of whether LH stimulation of granulosa cell ingression may contribute to these changes in the follicular structure that enable ovulation.

**Summary sentence:** LH-induced ingression of LH receptor-expressing cells within the mural granulosa layer of the ovarian follicle is a new component in the complex sequence of structural changes that lead to ovulation.

## Introduction

Preovulatory mammalian follicles are comprised of many concentric layers of cells. Directly around the oocyte are cumulus cells, and outside of these are a fluid-filled antrum, mural granulosa cells, a basal lamina, and a theca layer including steroidogenic cells, fibroblasts, smooth muscle cells, and vasculature [1] (Figure 1A). Ovulation is triggered when luteinizing hormone (LH), which is released from the pituitary and delivered to the ovary through blood vessels in the theca layer, acts on its receptors in a subset of the outer mural granulosa cells [2–8]. LH binding to its receptors initiates a G-protein-coupled signaling cascade that induces meiotic resumption in the oocyte and changes in gene expression in the granulosa cells that lead to ovulation [9–14].

**Figure 1.**
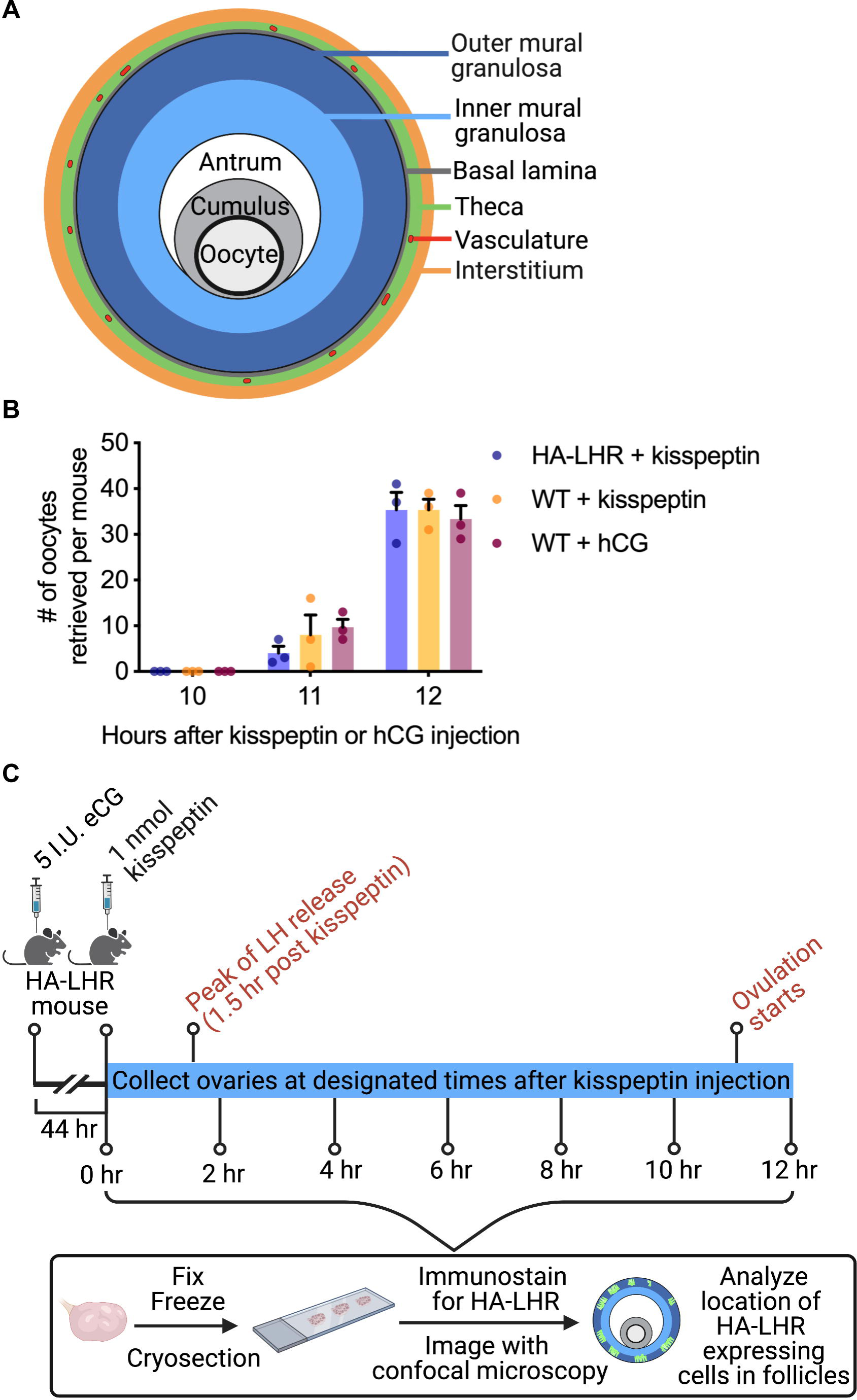
Follicle organization, time course of kisspeptin-induced ovulation, and experimental design. A) Tissue layers of a preovulatory mouse follicle. B) Time course of kisspeptin-induced ovulation in wild-type and HA-LHR mice. The time course of hCG-induced ovulation in wild-type mice is shown for comparison. Oviducts were collected at the indicated time points, and the numbers of ovulated oocytes were counted (n= 3 mice per each condition). C) Collection of ovaries after kisspeptin injection, for analysis of the localization of LH receptor-expressing cells by confocal microscopy.

Activation of the LH receptor also causes complex structural changes in the follicle. These include chromosomal and cytoskeletal rearrangements as meiosis progresses in the oocyte [15], secretion of an extracellular matrix from the cumulus cells that causes the cumulus mass to expand [16], and loss of connections between these cells and the oocyte as transzonal projections retract [17]. As ovulation approaches, the mural granulosa and theca layers in the apical region of the follicle, directly adjacent to the surface epithelium where the oocyte will be released, become thinner [18–21], correlated with vasoconstriction and reduced blood flow in the apical region of the theca [21]. The mural granulosa and theca layers in the basolateral region (that outside of the apical region) develop constrictions, and these are thought to contribute to expelling the oocyte and surrounding cumulus cells [22–27]. The preovulatory constrictions may be mediated in part by contraction of smooth muscle cells in the overlying theca [23], in response to endothelin-2 produced by the granulosa cells [26,28,29]. However, granulosa-specific deletion of endothelin-2 in mice only partially inhibits ovulation [30], suggesting that additional factors may also contribute to the follicular constriction. At the time of ovulation, the basal lamina around the granulosa cells breaks down, and blood vessels from the theca grow inwards as the remnants of the follicle transform into the corpus luteum, which produces progesterone to support pregnancy [31].

One unexplored factor that could contribute to the preovulatory changes in follicle structure is suggested by evidence that LH receptor activation induces migration [32,33] and cytoskeletal shape changes [34,35] in isolated mural granulosa cells. These cytoskeletal shape changes are detected within 30 minutes after LH receptor activation [35].

However, LH-stimulated granulosa cell motility has not been investigated in intact ovarian follicles. By using mice with an HA epitope tag on the endogenous LH receptor such that its localization could be visualized [8], and by collecting ovaries for imaging at defined times after eliciting a physiological LH surge by injection of kisspeptin [36], we obtained a precise timecourse of changes in localization of mural granulosa cells expressing the LH receptor over the period leading to ovulation. We discovered that LH stimulates granulosa cells that express its receptor to extend inwards in the follicle, intercalating between other granulosa cells. This LH-stimulated granulosa cell motility may influence the structure of the follicle as it prepares for ovulation and transformation into a corpus luteum.

## Materials and Methods

### Mice

Protocols covering the maintenance and experimental use of mice were approved by the Institutional Animal Care Committee at the University of Connecticut Health Center. The mice were housed in a room in which lights were turned on at 6 AM and off at 4 PM. Generation, genotyping, and characterization of mice with an HA tag on the endogenous LH receptor (HA-LHR) have been previously described [8]; homozygotes were used for breeding pairs and for all experiments. The HA-LHR and wildtype mouse background was C57BL/6J. These mice have been deposited at the Mutant Mouse Resource and Research Centers repository at The Jackson Laboratory (Bar Harbor, ME); MMRRC #71301, C57BL/6J-Lhcgr^em1Laj^/Mmjax, JR#038420.

### Hormone injections of prepubertal mice for collection of ovaries and assessment of ovulation

Ovaries containing preovulatory follicles were obtained by intraperitoneal injection of 22–24-day-old mice with 5 I.U. equine chorionic gonadotropin (eCG) (ProSpec #HOR-272). 44 hours later, the mice were injected intraperitoneally with 1 nmol kisspeptin-54 (Cayman Chemical, #24477) to stimulate an endogenous LH surge [36]. Ovaries were dissected at indicated time points after kisspeptin injection.

To assess ovulation, cumulus-oocyte complexes were collected from oviducts dissected after hormone injections as described above. For Figure 1B, some mice were injected with human chorionic gonadotropin (hCG, 5 I.U., ProSpec #HOR-250) instead of kisspeptin. Cumulus-oocyte complexes were dissociated by pipetting and then counted.

### Collection of ovaries from proestrus adult mice before and after the LH surge

Vaginal cytology of 6-8 week old females was examined daily at approximately 10 AM to determine the stage of the estrous cycle [37]. Ovaries from proestrus mice were collected at either 12 noon (pre-LH) or 10 PM (post-LH).

### Preparation of ovary cryosections for immunofluorescence microscopy

Ovaries were frozen, fixed, cryosectioned, and labelled for immunofluorescence microscopy as previously described [8]. Sections were 10 µm thick. Antibody sources and concentrations are listed in Table S1. Equatorial sections of follicles, defined as sections including the oocyte, were used for imaging.

### Confocal and Airyscan imaging

Ovarian cryosections were imaged with a confocal microscope (LSM800 or LSM980, Carl Zeiss Microscopy). To image entire follicle cross-sections, a 20x/0.8 NA Plan-Apochromat objective was used. Small regions of follicles were imaged using a 63x/1.4 NA Plan-Apochromat objective with an Airyscan detector to enhance resolution. Images of optical sections were reconstructed using the 2-dimensional Airyscan processing at standard strength. Except as indicated, all images were captured using an interval space of 1 µm. Brightness and contrast were adjusted after analysis in Fiji software [38].

### Image analysis

Preovulatory follicles were identified in equatorial cryosections by the presence of LH receptor-expressing cells in their mural granulosa cells. To quantify the percentage of LH receptor-expressing cell bodies that were located in the inner half of the mural granulosa layer (Figure 3A,B), inner and outer halves were defined as previously described [8]. In brief, the width of the mural granulosa layer from antrum to basal lamina was measured at 8 radial positions for each follicle, and halfway points were marked. These points were then connected to define the boundary of the inner and outer mural. The basal lamina position was identified by locating the outer edge of the outermost layer of granulosa cells, or by labeling with an antibody against laminin gamma 1. LH receptor-expressing cell bodies were identified as DAPI-stained nuclei that were surrounded by HA labeling, and were counted in inner and outer mural granulosa cell regions using the Cell Counter Tool in Fiji [38]. Counts were performed on the series of 10 optical sections with 1 µm interval space. The percentage of LH receptor-expressing cells in the inner mural was calculated by dividing the number of LH receptor-expressing cells in the inner mural by the total number of LH receptor-expressing cells in the inner and outer mural regions. The percentage of mural granulosa cells that expressed the LH receptor (Figure 3C) was calculated by dividing the total number LH receptor-expressing cells by the total number of DAPI-stained nuclei.

Basal lamina invaginations were analyzed using images of equatorial sections in which the basal lamina was labelled with an antibody against laminin gamma 1 (Table S1). Invaginations were defined as regions where the basal lamina curved inwards to a depth of 5-100 µm, with a width of ≤150 µm. The depth of each invagination (Figure 5C) was determined by drawing a line connecting the 2 points on the basal lamina at which it curved inwards, and then measuring the distance between that line and the deepest point of the invagination. For counts of the number of invaginations per cross-section (Figure 5D), the apical region of the follicle was defined as the mural granulosa region adjacent to the surface epithelium. The basolateral region was defined as the rest of the mural granulosa. Both measurements and counts were performed on the series of 10 optical sections with 1 µm interval space.

To measure the width of the mural granulosa layer in apical and basal regions, the center of the follicle was located by measuring the height and width of each follicle. A line was then drawn from the base through the center of the follicle to the apex, and beginning and end points were marked as the middle of the base and apex. Two points were marked at 50 µm along the basal lamina to the left and right of each point, and the width of the mural layer was measured at each point (see right hand panels of Figure 6A). The averages of the three measurements on the apical and basal sides were used for comparisons (Figure 6B). Measurements were performed on maximum projections of a series of 10 optical sections spaced 1 µm apart.

### Western blotting

For measurement of HA-LHR protein content (Figure S9), ovaries were collected at designated time points and sonicated in 1% SDS with protease inhibitors [39]. Protein concentrations were determined with a BCA assay (Thermo Scientific, #23227) and 40 µg of total protein was loaded per lane. Western blots were probed with an antibody against the HA epitope, developed using a fluorescent secondary antibody (Table S1), and detected with an Odyssey imager (LICOR, Lincoln, NE). Blots were co-imaged with the Revert stain for total protein (LICOR), and HA-LHR fluorescence intensity was normalized to the Revert fluorescence intensity for each lane. Values were then normalized to that for the ovary without kisspeptin injection.

### Statistics and graphics

Analyses were conducted as indicated in the figure legends using Prism 9 (GraphPad Software, Inc, La Jolla, CA). Values in graphs are presented as mean ± standard error of the mean (SEM), and values indicated by different letters are significantly different (P < 0.05). Diagrams for the graphical abstract and for Figures 1A,C and 3A were generated using BioRender.com.

## Results and Discussion

Except as indicated, LH release was stimulated by intraperitoneal injection of kisspeptin into ∼25 day old mice that had been injected 44 hours previously with eCG to stimulate follicle growth and LH receptor expression [36]. Kisspeptin is a neuropeptide that causes gonadotropin releasing hormone to be secreted from the hypothalamus, which in turn causes release of LH into the bloodstream [40]. The rise in serum LH peaks ∼1.5 hours after kisspeptin injection and is comparable in amplitude and duration to the endogenous LH surge [36]. As previously described, kisspeptin injection causes ovulation [36]. We chose to use kisspeptin rather than the more standard induction of ovulation by injection of human chorionic gonadotropin (hCG), because kisspeptin causes release of mouse LH, closely mimicking the natural ovulatory stimulus. To determine the kinetics of kisspeptin-induced ovulation, cumulus-oocyte complexes in the oviduct were counted at defined time points. With the experimental conditions used here, ovulation occurred between 11 and 12 hours after kisspeptin or hCG injection, and similar numbers of oocytes were released by each stimulus (Figure 1B).

The localization of cells expressing the LH receptor was investigated using a recently developed mouse line with a hemagglutinin (HA) tag on the endogenous LH receptor (HA-LHR), to allow specific immunolocalization of the LH receptor protein [8]. The heterogenous expression of the LH receptor within the mural epithelium allows individual cells to be well visualized [8]. The timing and number of oocytes released in response to kisspeptin were similar comparing HA-LHR and wild-type mice (Figure 1B). Ovaries from HA-LHR mice were collected, fixed, and frozen, either before kisspeptin injection, or at 2 hour intervals afterwards (Figure 1C). Ovary cryosections were then labelled with an HA antibody, and the localization of HA-LHR expressing cells was analyzed.

### LH induces ingression of LH receptor-expressing granulosa cells within the preovulatory follicle

The mural granulosa region of a mouse preovulatory follicle is comprised of ∼5-15 layers of cells, located between the basal lamina and the fluid-filled antrum [8,41,42] (Figures 1A, 2A, S1). Previously, we showed that before LH receptor stimulation, almost all of the LH receptor-expressing cell bodies in the mural granulosa are located in its outer half, and within this region, expression is heterogeneous [8]. Many of the LH receptor-expressing granulosa cells have an elongated flask-like shape, with the cell bodies containing their nuclei located a few cell layers away from the basal lamina and long processes extending back to the basal lamina, forming a pseudostratified epithelium [8,18] (Figures 2A, S1, S10). In the present study, only ∼7% of the total LH receptor-expressing granulosa cell bodies were found in the inner half of the mural granulosa region before LH receptor stimulation (Figure 3A,B).

**Figure 2.**
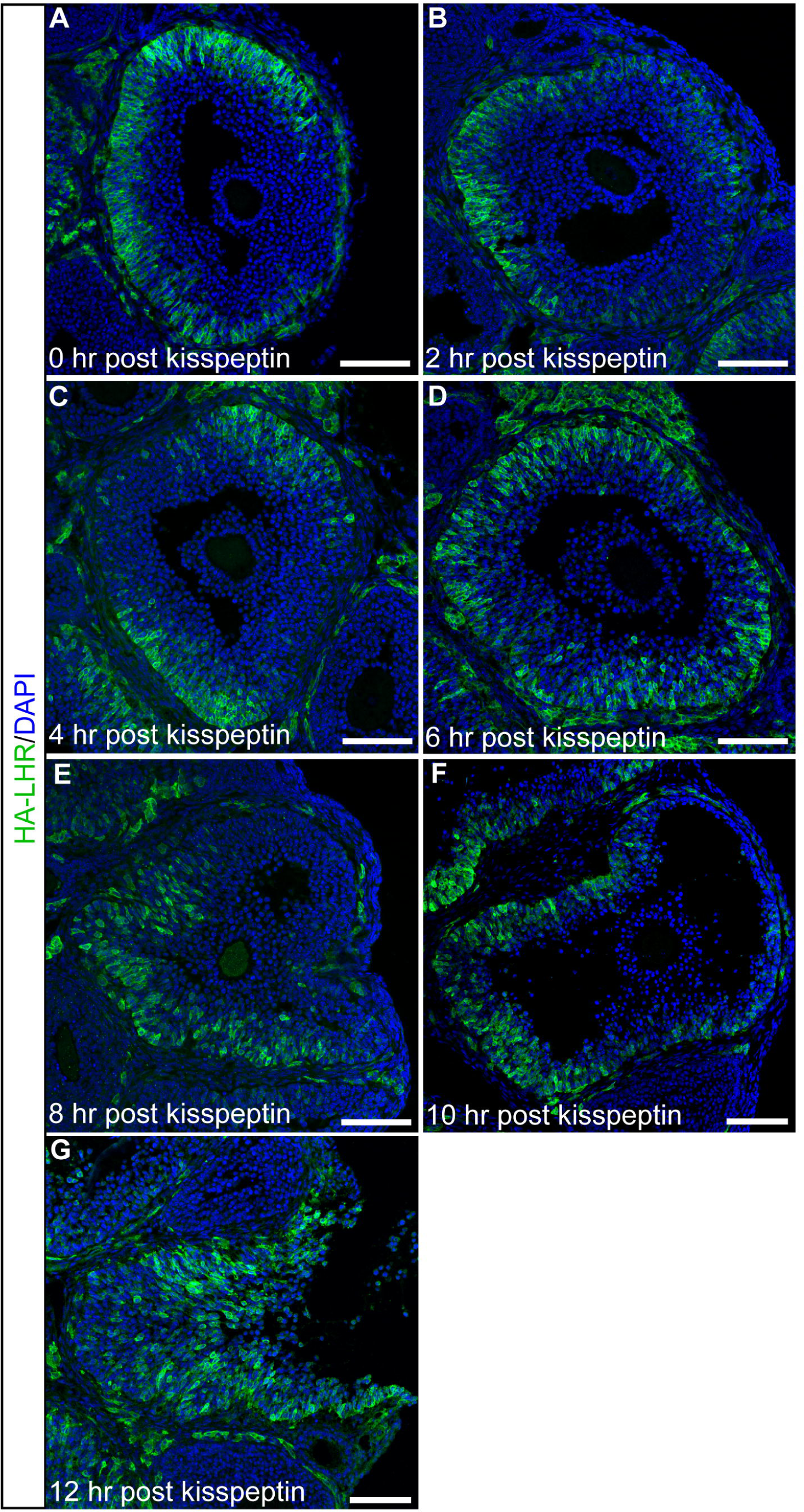
LH receptor-expressing cells extend inwards within the preovulatory follicle in response to LH. A-G) Confocal images of representative 10 µm thick equatorial cryosections of follicles in ovaries before and 2-12 hours after injection of HA-LHR mice with kisspeptin. The sections were labelled for HA immunofluorescence (green), and nuclei were labelled with DAPI (blue). Each panel shows a maximum projection of a stack of 10 optical sections imaged at 1 µm intervals with a 20x/0.8 NA objective. Scale bars = 100 µm. F and G were captured using a lower zoom than the other images to account for increased follicle size.

At 2 hours after kisspeptin injection (∼30 minutes after the peak of the induced LH surge), the percentage of LH receptor-expressing cell bodies in the inner mural layer had increased to ∼21% (Figures 2B, 3B, S2). At later time points after kisspeptin injection, this percentage continued to increase, reaching ∼35% at 10 hours (Figures 2C-F, 3B, S3-S6). Ovulation occurred between 11 and 12 hours after kisspeptin (Figures 1B, 2G, S7). Injection with PBS instead of kisspeptin did not change the percentage of LH receptor-expressing cells in the inner mural granulosa layer at 6 hours, indicating that the localization change is dependent on LH (Figures 3B, S8).

**Figure 3.**
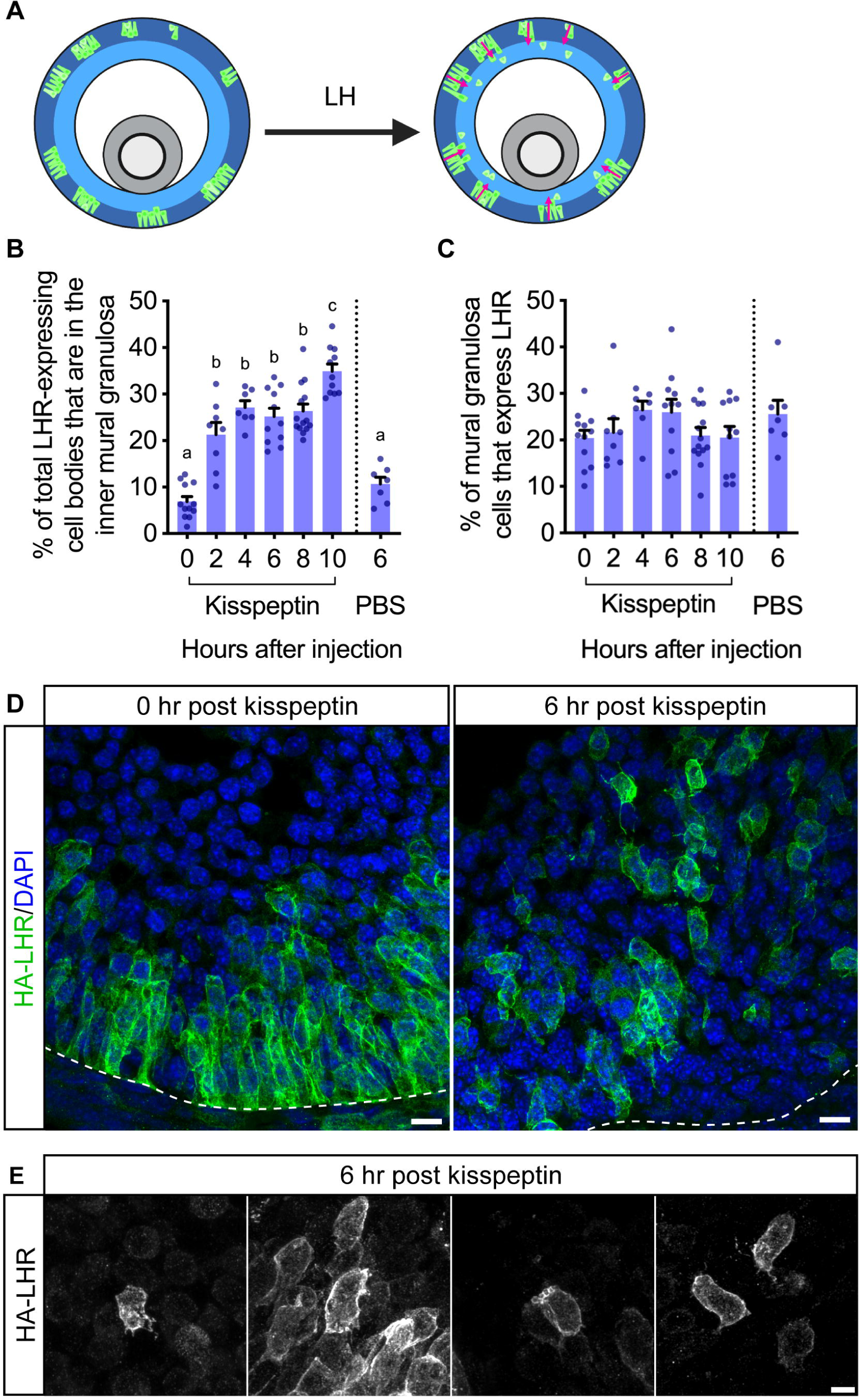
LH induces ingression of LH receptor-expressing cells and an epithelial-mesenchymal-like transition. A) Schematic of ingression of LH receptor-expressing cells within the follicle. B) Time course of the increase in the percentage of LH receptor (LHR)-expressing cell bodies in the inner half of the mural granulosa region after kisspeptin injection. No change in the distribution of LH receptor-expressing cells was seen at 6 hours after a control injection of PBS. C) No effect of kisspeptin on the percentage of mural granulosa cells that express the LH receptor. Measurements in B and C were made from images in Figures S1-S6 and S8. For each time point after kisspeptin, 1100-3000 cells from each of 7-14 follicles from 3-7 mice were analyzed. Each point on the graphs represents an individual follicle. Data were analyzed via one-way ANOVA with the Holm-Sidak correction for multiple comparisons. There were no significant differences among time points for data in C. D) Images of representative 10 µm thick equatorial cryosections of preovulatory follicles in ovaries from LH receptor-expressing mice before or 6 hours after kisspeptin injection. Each panel shows a maximum projection of a stack of 10 x 1 µm optical sections taken with a 63x/1.4 NA objective using the Airyscan detector. Scale bars = 10 µm. E) Filopodia and blebbing observed at the 6-hour time point. Images are maximum projections of a stack of 30 x 0.69 µm optical sections taken with 63x/1.4 NA objective using the Airyscan detector. Scale bar = 5 µm.

The increase in the fraction of LH receptor-expressing cells in the inner vs. outer mural granulosa region in response to the LH surge is most easily explained by migration into the inner region of cells originally in the outer region. This conclusion is consistent with previous studies showing that LH receptor stimulation causes granulosa cells in culture to become migratory [32,33]. Future live imaging of granulosa cell migration in intact follicles, using mice in which cells expressing the LH receptor are fluorescently tagged, could further test this conclusion.

Alternatively, the observed increase in the ratio of LH receptors in the inner vs. outer mural granulosa regions could potentially be a consequence of synthesis of LH receptor mRNA and protein in the inner mural granulosa region. Our previously published in situ hybridization data showed no detectable LHR mRNA in the inner mural granulosa cells in follicles prior to the LH surge [8], so such an explanation would require new transcription of LHR mRNA. In rat ovaries, LHR mRNA synthesis has not been detected until 24-72 hours after hCG injection [6,43], making this an unlikely explanation of the relocalization of LHR protein that we observe in mouse preovulatory follicles beginning within ∼30 minutes after the peak of the LH surge (2 hours after kisspeptin injection).

Nevertheless, to examine this alternative hypothesis, we compared the percentage of the total population of mural granulosa cells that expressed LH receptor protein over the 10-hour period after kisspeptin injection. No significant increase was seen over this time period, and mean values at 6 hours after injection of kisspeptin or a control solution of PBS were identical (Figure 3C). These results are consistent with the conclusion that the LH surge did not stimulate synthesis of its receptor over this time period. We also measured the LH receptor protein content of ovaries after kisspeptin injection, using quantitative western blotting, and found that LH receptor protein levels were unchanged over the 12-hour period after injection (Figure S9). If new LHR protein synthesis occurred in the inner mural region, we would expect to have seen an increase in the total LH receptor content of the ovary, unless an equal amount of LH receptor protein in the outer mural region was proteolyzed at the same time. Thus, new synthesis of LH receptor protein in the inner mural granulosa layer is an unlikely explanation of the observed redistribution of LH receptor-expressing cells in the follicle.

### LH induces an epithelial-to-mesenchymal-like transition in LH receptor-expressing granulosa cells within the follicle

High resolution Airyscan images of ovaries from mice before and after kisspeptin injection showed that by 6 hours after kisspeptin injection, many LH receptor-expressing cells in the follicle interior were rounder compared to the predominantly flask-shaped cells seen near the basal lamina prior to LH receptor stimulation (Figures 3D, S10, S11). At 6 hours, many of the LH receptor-expressing cells had lost a visible attachment to the basal lamina, suggesting that these cells had detached, undergoing an epithelial-to-mesenchymal-like transition (Figures 3D,E, S11). Alternatively, the cellular processes connecting these cell bodies to the basal lamina might have been too thin to see with light microscopy or might not have been contained within the 10 µm thick section. These cells often had numerous filopodia extending in many directions, as well as membrane blebs (Figure 3D,E), consistent with the presence of filopodia and blebs on other migratory cells [44–46].

### The LH surge also induces ingression of LH receptor-expressing cells in preovulatory follicles of naturally cycling adult mice

To confirm that the ingression of granulosa cells that was seen in hormonally stimulated prepubertal mice also occurred in naturally cycling mice, we collected ovaries from adult mice in proestrus. Based on measurements of serum LH, the peak of the proestrous LH surge occurs close to the time that the lights are turned off in the room where the mice are housed, although the exact time is variable [47]. Mice were determined to be in proestrus based on vaginal cytology, and we collected ovaries at either 4 hours before lights off (pre-LH) or 6 hours after lights off (post-LH). In the pre-LH ovaries, ∼11% of LH receptor expressing cells were located in the inner mural, whereas in the post-LH ovaries, the LH receptor-expressing cell bodies had moved inwards, with ∼35% of cells expressing the LH receptor localized in the inner mural (Figures 4A,B, S12). This was comparable to the change in localization that occurred in the prepubertal mice (Figures 2, 3B), confirming that the ingression occurs during the natural ovulation process.

**Figure 4.**
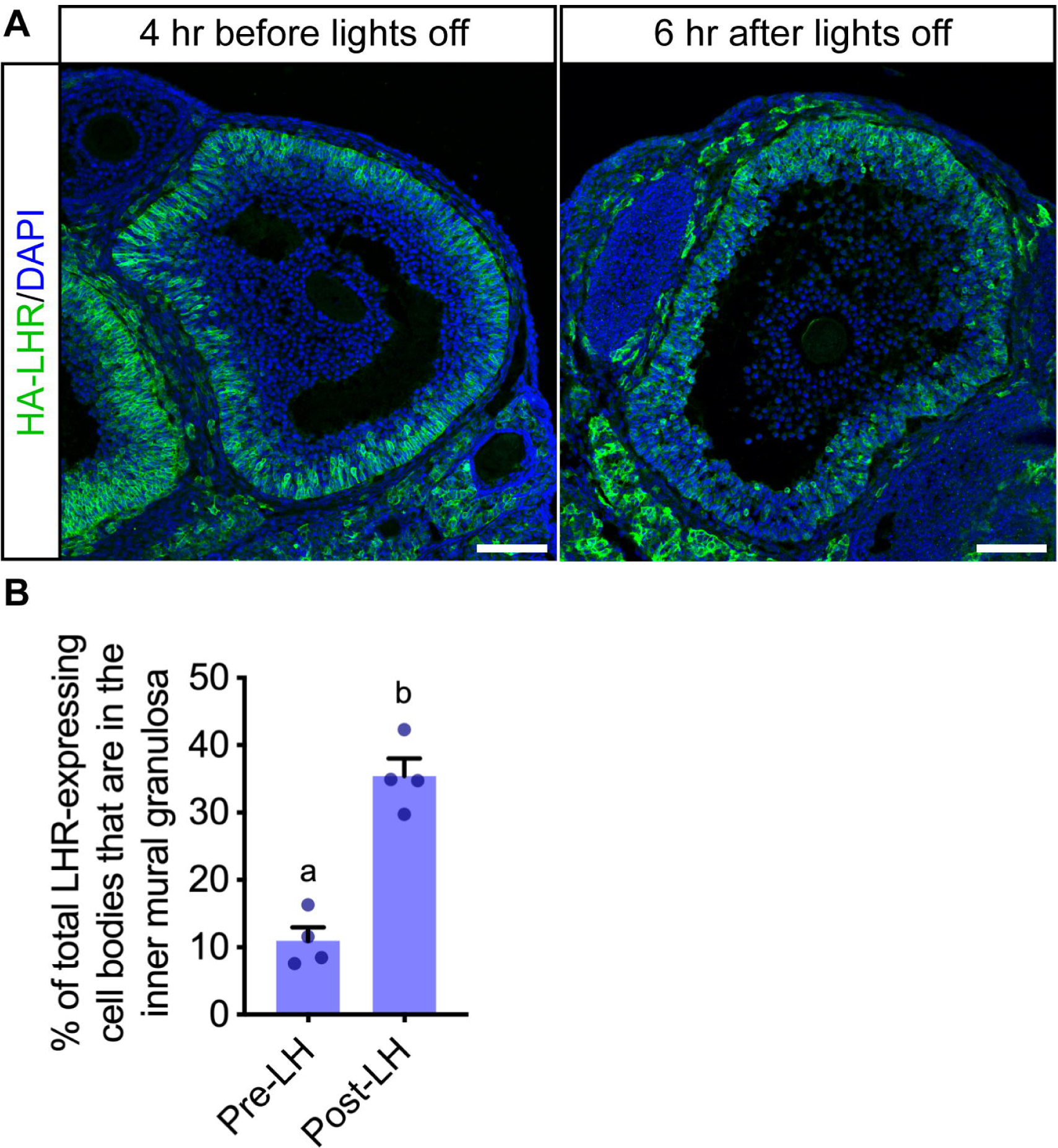
LH induces ingression of LH receptor-expressing cells in follicles of naturally cycling mice. A) Representative images of follicles from mice collected either 4 hours before lights off (pre-LH) or 6 hours after lights off (post-LH) on the day of proestrus. The sections were labelled for HA immunofluorescence (green), and nuclei were labelled with DAPI (blue). Each panel shows a maximum projection of a stack of 10 optical sections imaged at 1 µm intervals with a 20x/0.8 NA objective. Scale bars = 100 µm. Note that the section through the edge of the follicle at the far left of panel A was not equatorial, accounting for the difference in appearance. B) The percentage of LH receptor-expressing cell bodies in the inner half of the mural granulosa region in mice before or after the endogenous LH surge. Measurements were made from follicles in Figure S12. Each point on the graphs represents an individual follicle; data were collected from 2 mice at each time point. Data were analyzed via an unpaired t-test (p < 0.0005).

### LH induces invaginations of the basolateral surface of the follicle, starting ∼6 hours before ovulation

The LH-induced ingression of the LH receptor-expressing granulosa cells into the inner half of the mural layer, and the associated changes in cell shape, raised the question of how these cellular events might correlate with LH-induced changes in the shape of the follicle as a whole. Constrictions in the basolateral surface of follicles have been previously observed during or within 2 hours of ovulation, but reports at earlier time points are lacking [24–29]. By fixing ovaries before or 6-10 hours after injection of mice with kisspeptin, labelling the basal lamina in ovary cryosections, and imaging equatorial cross-sections of preovulatory follicles, we observed that invaginations of the follicle surface began much earlier than previously reported.

In follicles in ovaries from mice that had not been injected with kisspeptin, the basal lamina was smooth, with only occasional inward deflections (Figure 5A,B). However, at 6-10 hours after kisspeptin injection, the basal lamina showed numerous invaginations, defined as regions where the basal lamina curved inwards to a depth of 5-100 µm with a width of ≤150 µm (Figure 5A-C). The invaginations occurred only in basolateral regions of the follicle, and were seen as early as 6 hours after kisspeptin injection, corresponding to ∼4.5 hours after the peak of the LH surge, and ∼6 hours before ovulation (Figures 5A-D, S4B, S5B, S6B). Their number increased between 6 and 10 hours after kisspeptin injection (Figure 5D). In addition to the invaginations, the basolateral region of follicles at 8 and 10 hours after kisspeptin injection usually also showed larger scale constrictions (Figures 5B, S5, S6). Similar invaginations and constrictions of the basolateral surface were seen in follicles of ovaries after the LH surge in naturally cycling mice (Figures 4A, S12).

**Figure 5.**
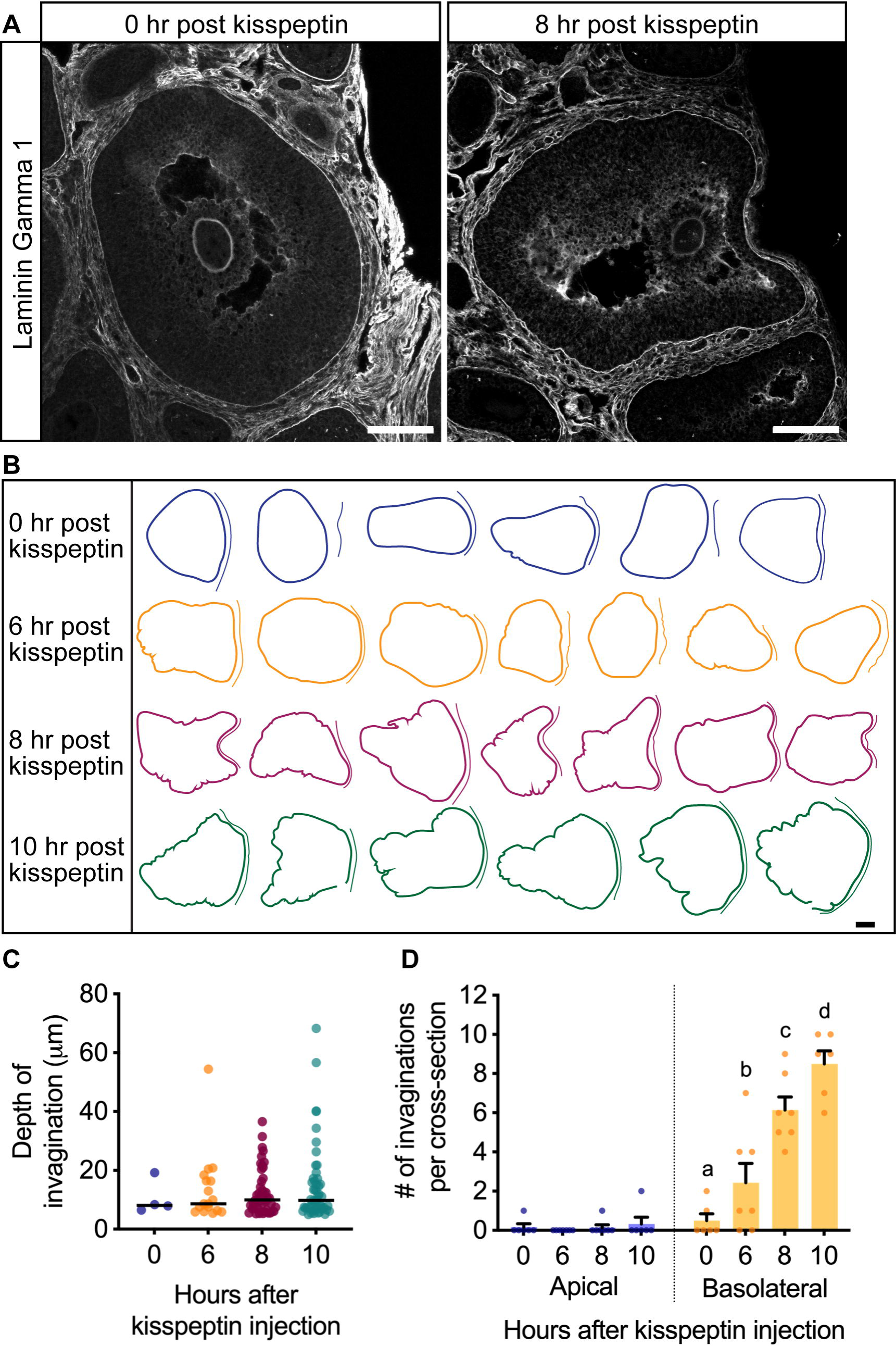
LH induces basal lamina invaginations and constrictions in the basolateral region of the follicle. A) Representative images of follicles before kisspeptin injection and 8 hours afterwards, labeled for laminin gamma 1 (white). Scale bars = 100 µm. B) Tracings of the basal lamina of follicles in equatorial sections at 0, 6, 8, or 10 hours after kisspeptin injection (from images in Figures S1B, S4B, S5B, S6B). The tracings are aligned such that the apical region is on the right, as indicated by the double line. Apical was defined as the region that is directly adjacent to the surface epithelium, while basolateral was defined as any portion of the follicle that is not in contact with the surface epithelium. Scale bar = 100 µm. C) Depth of invaginations in follicles at 0, 6, 8, and 10 hr post kisspeptin. Each symbol represents one invagination. D) Total number of invaginations in apical and basolateral regions of follicles at 0, 6, 8, and 10 hr post kisspeptin. Each symbol represents one follicle. Data for C and D were generated using datasets in Figures S1B, S4B, S5B, S6B). Each point on the graphs represents an individual follicle; data were collected from 6 - 7 mice at each time point. Data were analyzed via two-way ANOVA with the Holm-Sidak correction for multiple comparisons.

### LH induces thinning of the apical surface of the follicle, starting ∼6 hours before ovulation

In addition to the basolateral constrictions of the follicle surface, LH stimulation caused thinning of the mural granulosa layer at the apex of the follicle where the oocyte will be released (Figures 2, 5A, 6, S4-S7). As previously reported for hamster follicles [20], the apical thinning begins early, showing a statistically significant decrease in mean thickness by 6 hours after injection of kisspeptin, and not occurring in the basal region (Figure 6B). At 8 hours after kisspeptin, many follicles also had a concavity on the apical side, which was wider and smoother than the invaginations seen in basolateral regions (Figures 5A,B, S5). Apical thinning was evident in some of the follicles of naturally cycling mice after the LH surge; variability in the timing of the LH surge with respect to the time of lights off may explain why it was not seen in all examples (Figures 4A, S12).

**Figure 6.**
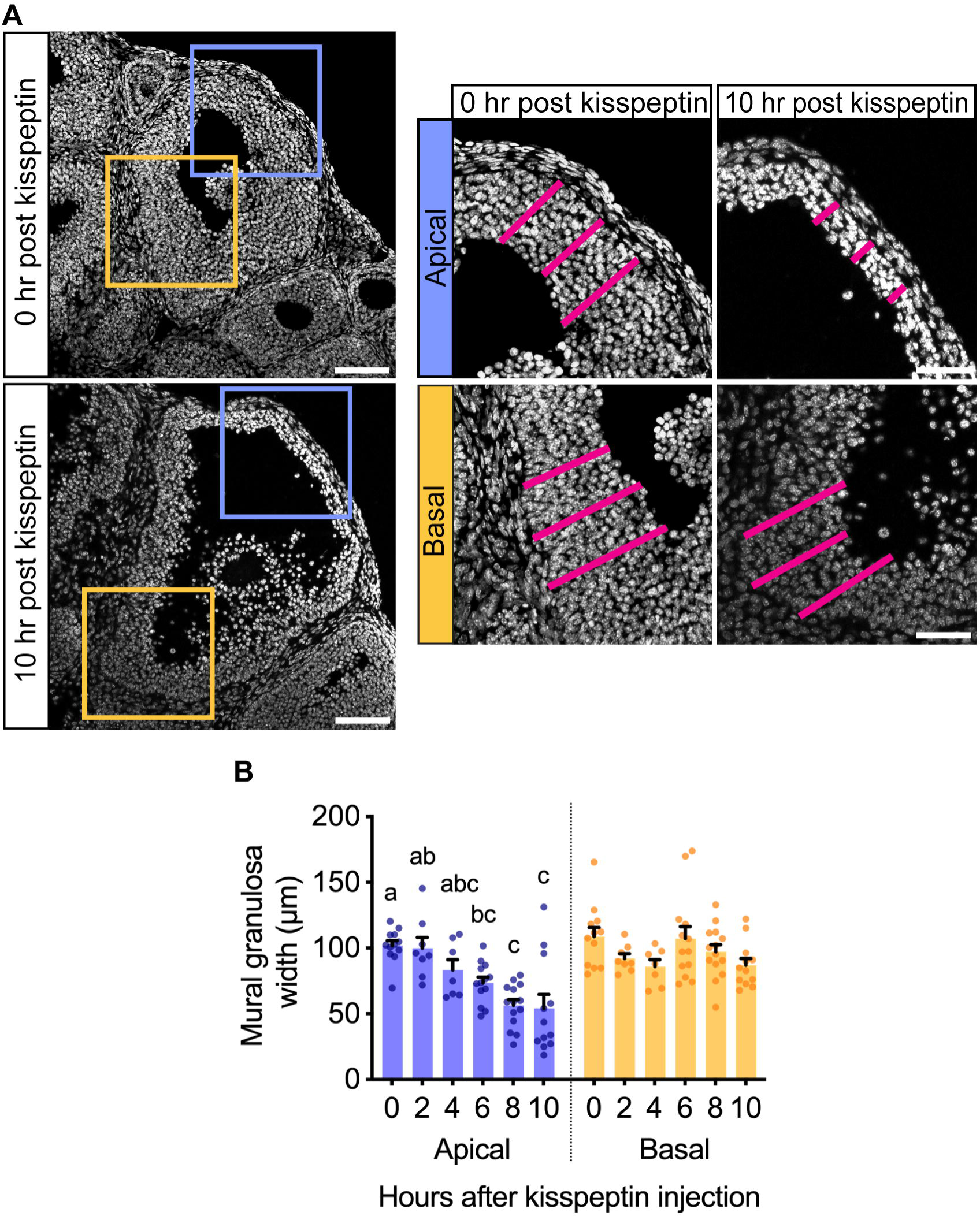
LH induces thinning of the mural granulosa layer at the apex but not the base. A) Representative images of follicles before kisspeptin injection and 10 hours after, with nuclei labelled with DAPI (white). Apical and basal regions (blue and yellow boxes) are enlarged at the right. Pink lines represent measurements of the mural granulosa width, taken 50 µm apart and then averaged to generate the points in (B). Scale bars = 100 µm for full follicle images on the left, 50 µm for insets on the right. B) Width of mural granulosa layer in apical and basal regions of follicles before and 2-10 hours after kisspeptin injection. Each point on the graph represents the average measurements from one follicle from Figures S1-S6. Data were analyzed via one-way ANOVA with the Holm-Sidak correction for multiple comparisons.

### Does LH-induced ingression of LH receptor-expressing mural granulosa cells contribute to LH-induced basolateral invaginations or apical thinning?

Our findings that basolateral invaginations and apical thinning are both evident by 6 hours after kisspeptin injection (Figures 5 and 6) raise the question of whether the ingression of LH receptor-expressing cells might contribute to causing these events. Inwardly extending granulosa cells could be speculated to pull the attached basal lamina inwards, contributing to LH-induced invaginations and constrictions of the basolateral follicle surface. Likewise, migration of granulosa cells out of the apical region of the mural granulosa could be imagined to contribute to thinning of the follicle apex. While previous studies have provided evidence for the combined action of extracellular proteases [13,21,27] and endothelin-induced contractions of smooth muscle cells in the theca [29] and theca blood vessels [21] in causing both basolateral invaginations and apical thinning, definitive identification of the cells that are responsible remains uncertain.

We hypothesize that LH-induced ingression of LH receptor-expressing mural granulosa cells may be an additional factor that contributes to LH-induced changes in follicle shape. In support of this concept, cell migration has been found to regulate tissue architecture in other developmental systems [48,49], including *Drosophila* ovarian follicles [50] and the mammalian placenta [51]. LH-stimulated granulosa cell migration could provide a complementary mechanism that acts in parallel with proteases [13,21,27], and with contraction of vascular and non-vascular smooth muscle cells in the theca layer [21,23,24,28–30] to contribute to rupture of the follicle surface at ovulation.

Importantly, the early changes in follicular shape reported here start around 6 hours after kisspeptin injection, apparently preceding the synthesis of endothelin mRNA by the granulosa cells, which has been reported to begin at ∼11 hours after injection of mice with the LH receptor agonist hCG [28]. Based on these kinetics, endothelin-induced responses do not appear to fully account for the LH-induced changes constrictions in the basolateral region of the follicle and the thinning of the mural granulosa layer in the apical region, although additional spatial-temporal analysis of the expression of endothelins and their receptors would be useful.

Understanding of the possible contribution of LH-induced ingression of mural granulosa cells to causing the invaginations in the basolateral region of the follicle and the thinning of the apical mural granulosa cell layer could be furthered by knowledge of the cytoskeletal changes that cause the cells to migrate, such that the migration could be inhibited. Notably, activation of the actin severing and depolymerizing protein cofilin is required for LH receptor-stimulated granulosa cell shape changes in vitro [35], suggesting a possible function of cofilin in granulosa cell ingression within the follicle in vivo. Phosphoproteomic analysis of rat follicles has shown that LH signaling decreases the phosphorylation of cofilin on serine 3 to ∼10% of baseline within 30 minutes [52], and dephosphorylation at this site increases cofilin activity [53]. Cofilin is also essential for other developmental processes involving cellular extension, such as axon growth [54]. Mice with genetically modified cofilin or other cytoskeletal proteins in their granulosa cells could be used to investigate the mechanisms that mediate LH-induced granulosa cell ingression in preovulatory follicles, and the consequences for follicular shape changes and ovulation. Happening over a similar time frame as LH-induced changes in follicle shape, LH-induced ingression of LH receptor-expressing cells within the mural granulosa layer is a new component in the complex sequence of structural changes in the follicle that occur in the hours preceding ovulation.

## Supplementary material

Table S1. Antibodies used for this study.

Figures S1-S8. Images of follicles in ovaries from mice 0-12 hours after kisspeptin injection or 6 hours after PBS injection.

Figure S9. Quantitative western blot analysis of HA-LHR protein in ovaries from mice 0-12 hours after kisspeptin injection or 6 hours after PBS injection.

Figure S10-S11. High resolution images of granulosa cells within follicles in ovaries without kisspeptin injection or 6 hours after injection.

Figure S12. Images of follicles in ovaries from adult proestrus mice, collected at either 4 hours before or 6 hours after lights were turned off.

## Data availability

All relevant data can be found within the article and its supplementary information.

## Author contributions

CMO and LAJ designed and carried out the studies, analyzed the data, and wrote the manuscript.

## Conflict of interest

The authors declare that no conflict of interest exists.

## Acknowledgements

We thank Deb Kaback and Tracy Uliasz for technical assistance, Lydia Sorokin and Siegmund Budny for generously providing the laminin-gamma 1 antibody, Mark Terasaki, Jeremy Egbert, Rachael Norris, Siu-Pok Yee, Lisa Mehlmann, Valentina Baena, Allan Herbison, Tabea Marx, Chris Thomas, and Melina Schuh for helpful discussions, and John Eppig for critical reading of the manuscript.

## Abbreviations

LH: luteinizing hormone
LHR: luteinizing hormone receptor
HA-LHR: hemagglutinin-tagged luteinizing hormone receptor
eCG: equine chorionic gonadotropin
hCG: human chorionic gonadotropin

## Grant Support

This work was supported by the *Eunice Kennedy Shriver* National Institute of Child Health and Human Development (R37 HD014939 to L.A.J., and F31 HD107918 to C.M.O.).

---

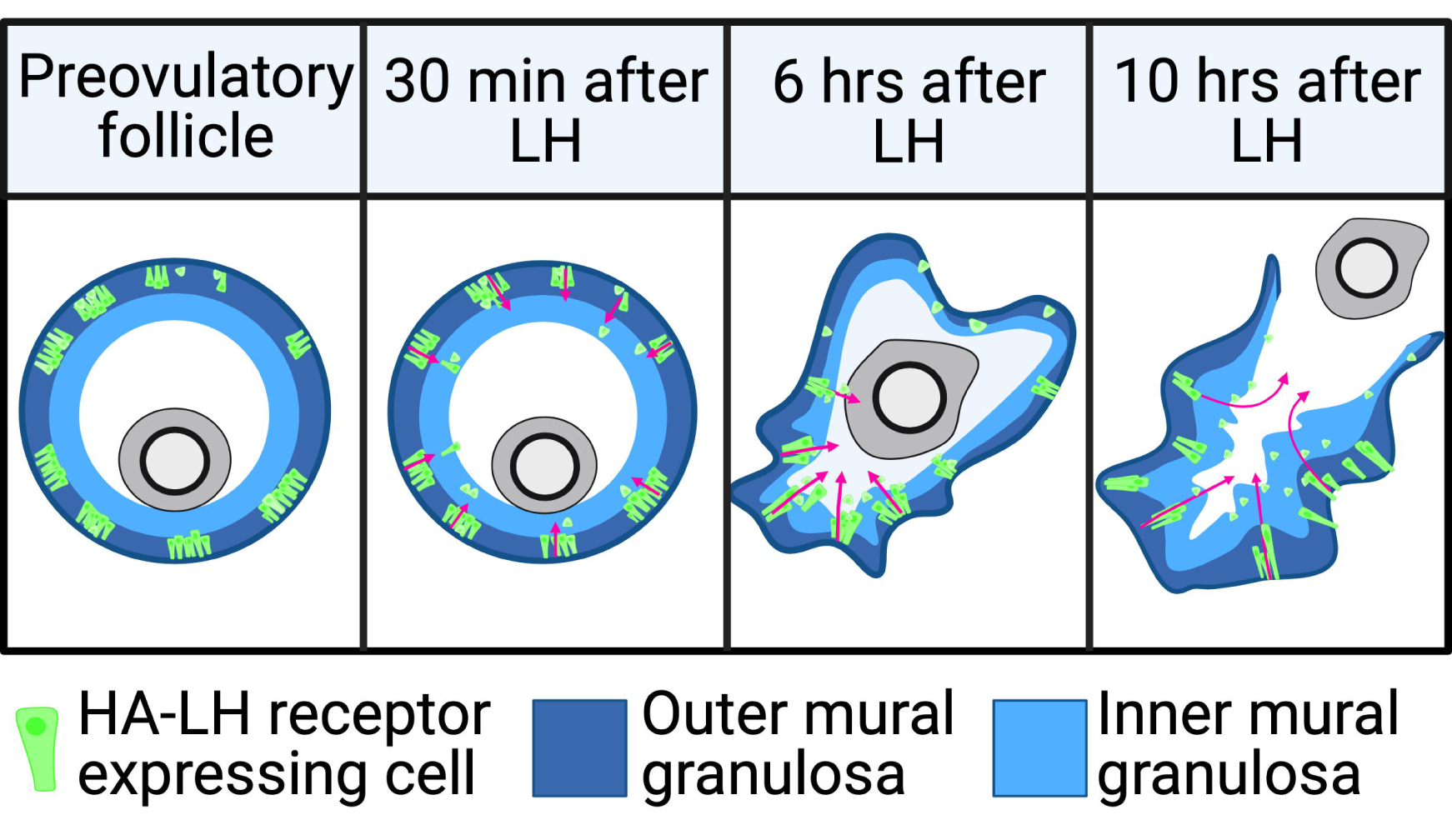

## Notes

### Competing Interest Statement

The authors have declared no competing interest.

### Summary of Updates

Text updated to clarify results and discussion; figure 3 and 4 updated

