## Supplementary material for "Luteinizing hormone stimulates ingression of mural granulosa cells within the mouse preovulatory follicle": Table S1

**Table S1.** Antibodies used for this study.

| Antibody | Source | Final Concentration |
| --- | --- | --- |
| rabbit monoclonal antibody against HA-tag; RRID: AB_1549585 | Cell Signaling, 3724 | Immunofluorescence: 0.13 µg/mL Western blots: 0.067 µg/mL Stock: 67 µg/mL |
| rat anti-mouse laminin gamma 1 chain, clone 3E10; RRID: AB_1123687 | Provided by Lydia Sorokin, University of Muenster [55, 56] | Conditioned medium, undiluted |
| goat anti-rabbit IgG (H+L) Cross-Adsorbed Secondary Antibody, Alexa Fluor-488; RRID: AB_257627 | Thermo Fisher Scientific, Molecular Probes, A- 11034 | 4 µg/mL |
| goat anti-rat IgG (H+L) Cross-Adsorbed Secondary Antibody, Alexa Fluor-555; RRID: AB_2535855 | Thermo Fisher Scientific, Molecular Probes, A- 21434 | 4 µg/mL |
| IRDye 800CW goat anti- Rabbit IgG Secondary Antibody; RRID: AB_621843 | LI-COR, 926-32211 | 0.067 µg/mL |
