## Supplementary material for "Luteinizing hormone stimulates ingression of mural granulosa cells within the mouse preovulatory follicle": Figure S1-12

**Figure S1.** Images of follicles from mice without kisspeptin injection, used to generate data in Figures 3B, 3C, and 6B. A) Follicles labeled for HA-LHR and DAPI. B) Follicles labeled for HA-LHR, DAPI and laminin gamma 1. Images in Figure S1B were also used to generate data in Figures 5C and 5D. Scale bars = 100 µm.

**Figure S2.** Images of follicles 2 hours after kisspeptin injection, labeled for HA-LHR and DAPI and used to generate data in Figures 3B, 3C, and 6B. Scale bars = 100 µm.

**Figure S3.** Images of follicles 4 hours after kisspeptin injection, labeled for HA-LHR and DAPI and used to generate data in Figures 3B, 3C, and 6B. Scale bars = 100 µm.

**Figure S4.** Images of follicles 6 hours after kisspeptin injection, used to generate data in Figures 3B, 3C, and 6B. A) Follicles labeled for HA-LHR and DAPI. B) Follicles labeled for HA-LHR, DAPI, and laminin gamma 1. Images in Figure S4B were used to generate data in Figures 5C and 5D. Scale bars = 100 µm.

**Figure S5.** Images of follicles 8 hours after kisspeptin injection, used to generate data in Figures 3B, 3C, and 6B. A) Follicles labeled for HA-LHR and DAPI. B) Follicles labeled for HA-LHR, DAPI, and laminin gamma 1. Images in Figure S5B were used to generate data in Figures 5C and 5D. Scale bars = 100 µm.

**Figure S6.** Images of follicles 10 hours after kisspeptin injection, used to generate data in Figures 3B, 3C, and 6B. A) Follicles labeled for HA-LHR and DAPI. B) Follicles labeled for HA-LHR, DAPI, and laminin gamma 1. Images in Figure S6B were used to generate data in Figures 5C and 5D. Scale bars = 100 µm.

**Figure S7.** Images of follicles 12 hours after kisspeptin injection. Scale bars = 100 µm. White dotted lines represent oocytes in the process of being ovulated.

**Figure S8.** Images of follicles 6 hours after PBS injection, labeled for HA-LHR and DAPI, and used to generate data in Figures 3B and 3C. Scale bars = 100 µm.

**Figure S9.** Quantitative western blot analysis of HA-LHR protein in 40 µg of total ovary protein from HA-LHR mice, before and 4, 8, 10, and 12 hours after injection of kisspeptin. The WT lane (40 µg of wildtype ovary) shows the specificity of the HA antibody. The predicted MW for the HA-LHR polypeptide is 79 kDa; the doublet band located above the 75 kDa molecular weight marker may represent differently glycosylated forms. Lower MW bands may represent breakdown products or splice variants. The graph shows analysis of three similar blots. Blots were co-imaged with the Revert stain for total protein (LICOR), and HA-LHR fluorescence intensity was normalized to the Revert fluorescence intensity for each lane. HA-LHR concentrations are expressed as a ratio of the amount of HA-LHR protein per total ovary protein, normalized to value of 1.0 A.U. for the ovary without kisspeptin injection. There were no significant differences among time points (one-way ANOVA).

Previous studies of rat ovaries have shown that LH receptor stimulation results in a decrease in LH receptor mRNA content [6,43,57-59] and LH receptor-ligand binding [2,43] during the initial 12-hour period after hCG injection. Thus, we were initially surprised that over the time course of our studies, LH receptor stimulation did not decrease the LH receptor content of the mouse ovary or the number of granulosa cells expressing the LH receptor (Figure 3C). Possible explanations for this apparent difference from previous reports include differences in hormonal stimulation protocols [59] and differences between rats and mice. In addition, it is possible that as previously noted [43], the observed decrease in LH receptor-ligand binding could be due to the internalization of the LH receptor in granulosa cells [12] rather than a decrease in total LH receptor protein.

**Figure S10.** High resolution images of granulosa cells from ovaries without kisspeptin injection, labeled for HA-LHR and DAPI. Scale bars = 10 µm.

**Figure S11.** High resolution images of granulosa cells from ovaries 6 hours after kisspeptin injection, labeled for HA-LHR and DAPI. Scale bars = 10 µm.

**Figure S12.** Images of follicles from adult mice either 4 hours before lights off (A) or 6 hours after lights off (B) on the day of proestrus, used to generate data in Figure 4B. A) Follicles labeled for HA-LHR and DAPI. Scale bars = 100 $\mu$m.

Figure S1

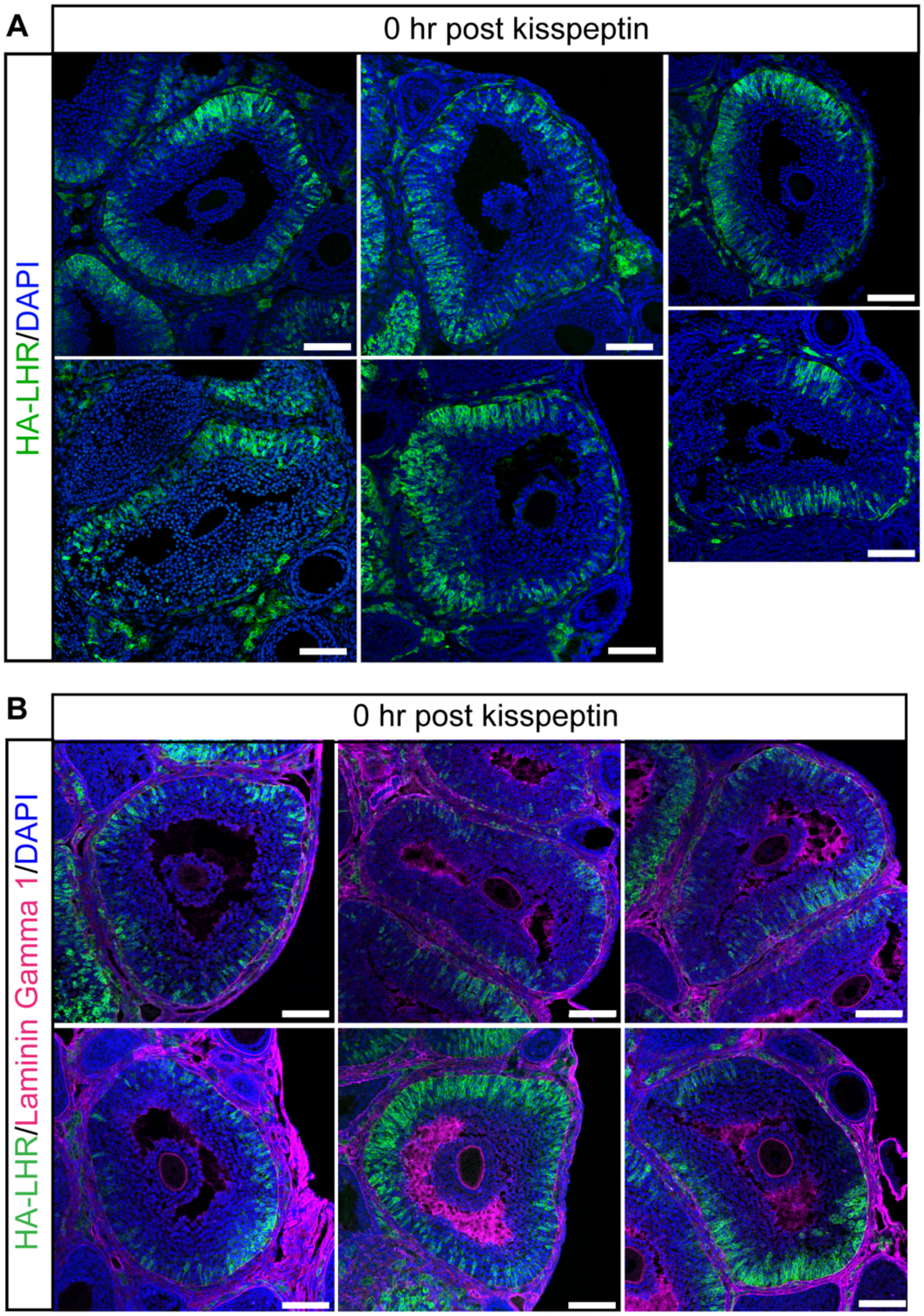
A
0 hr post kisspeptin
HA-LHR/DAPI
B
0 hr post kisspeptin
HA-LHR/Laminin Gamma 1/DAPI

Figure S2

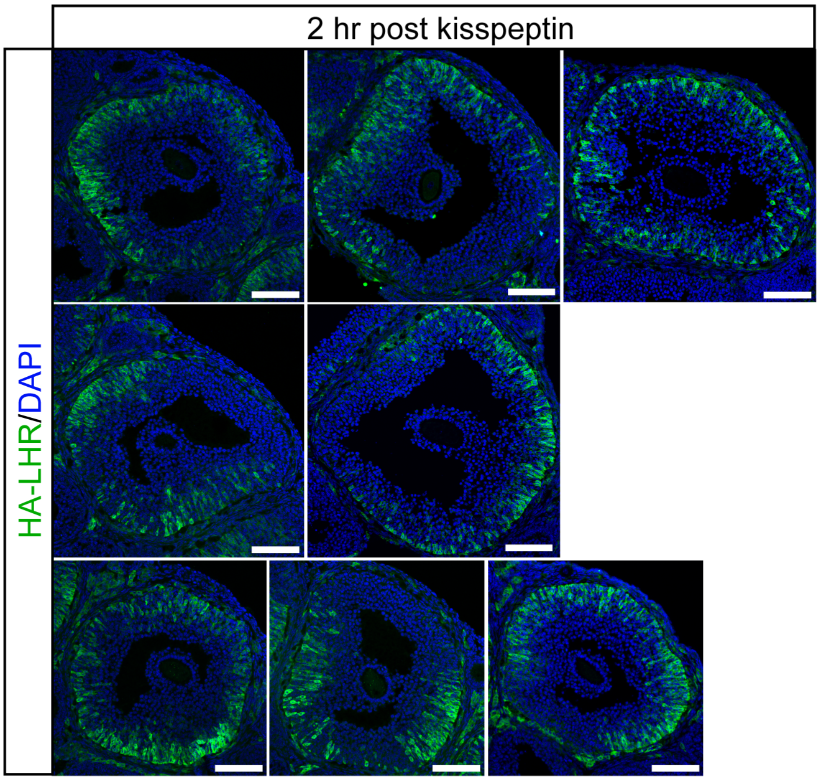

Figure S3

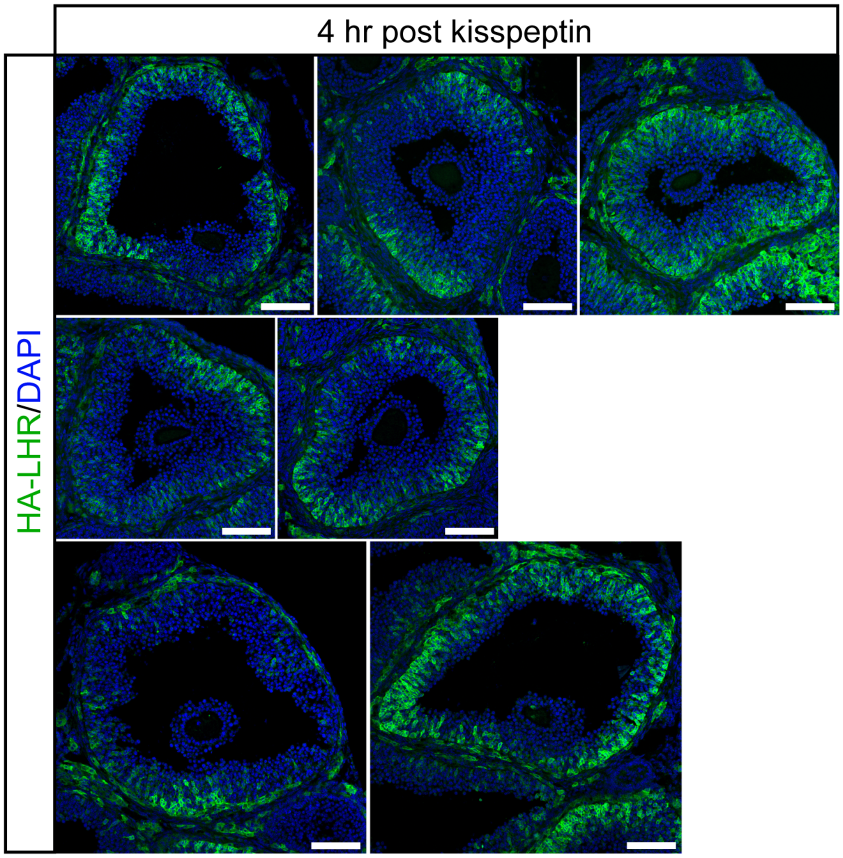

Figure S4

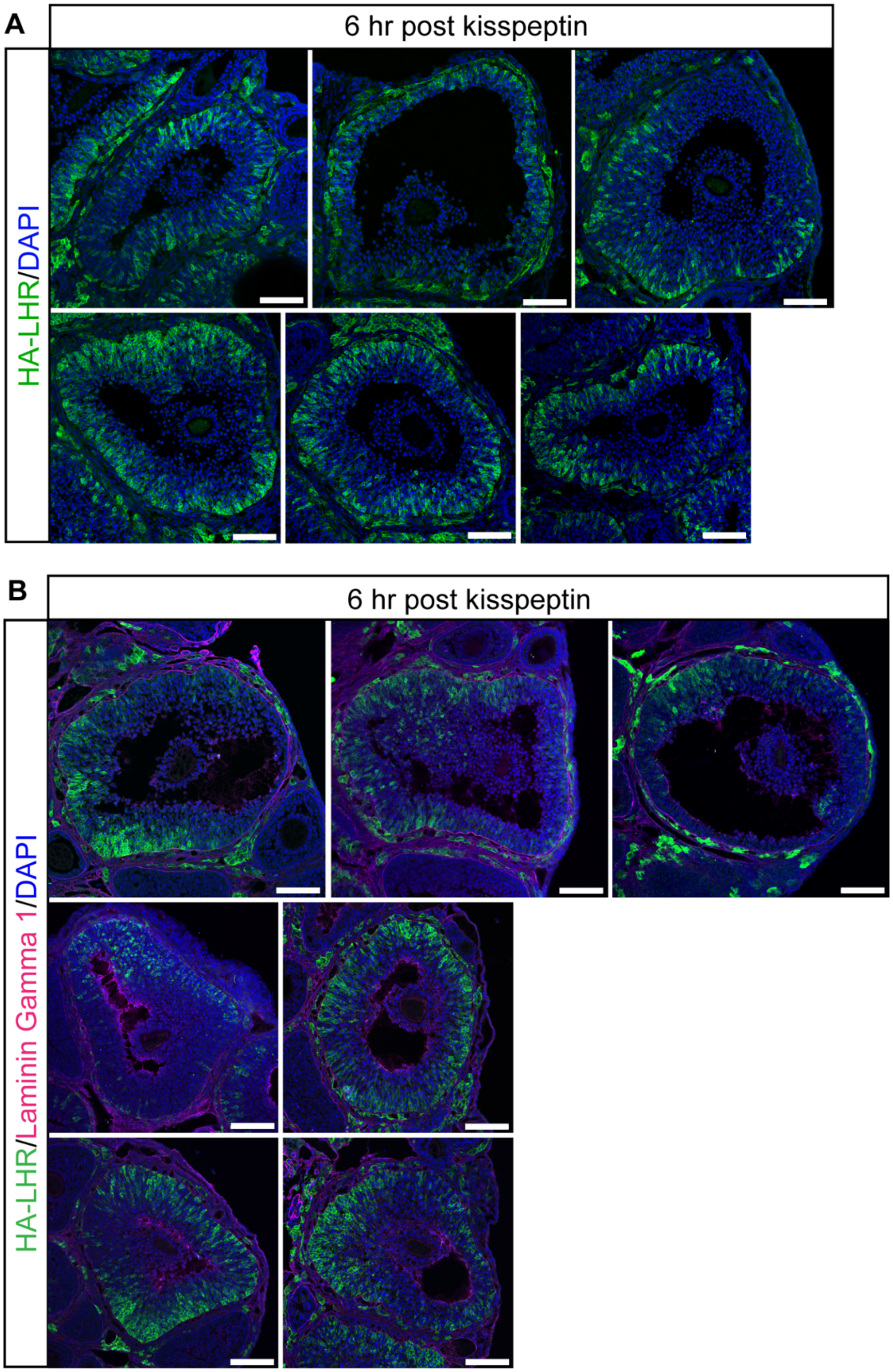
A
6 hr post kisspeptin
HA-LHR/DAPI
B
6 hr post kisspeptin
HA-LHR/Laminin Gamma 1/DAPI

116 Figure S5

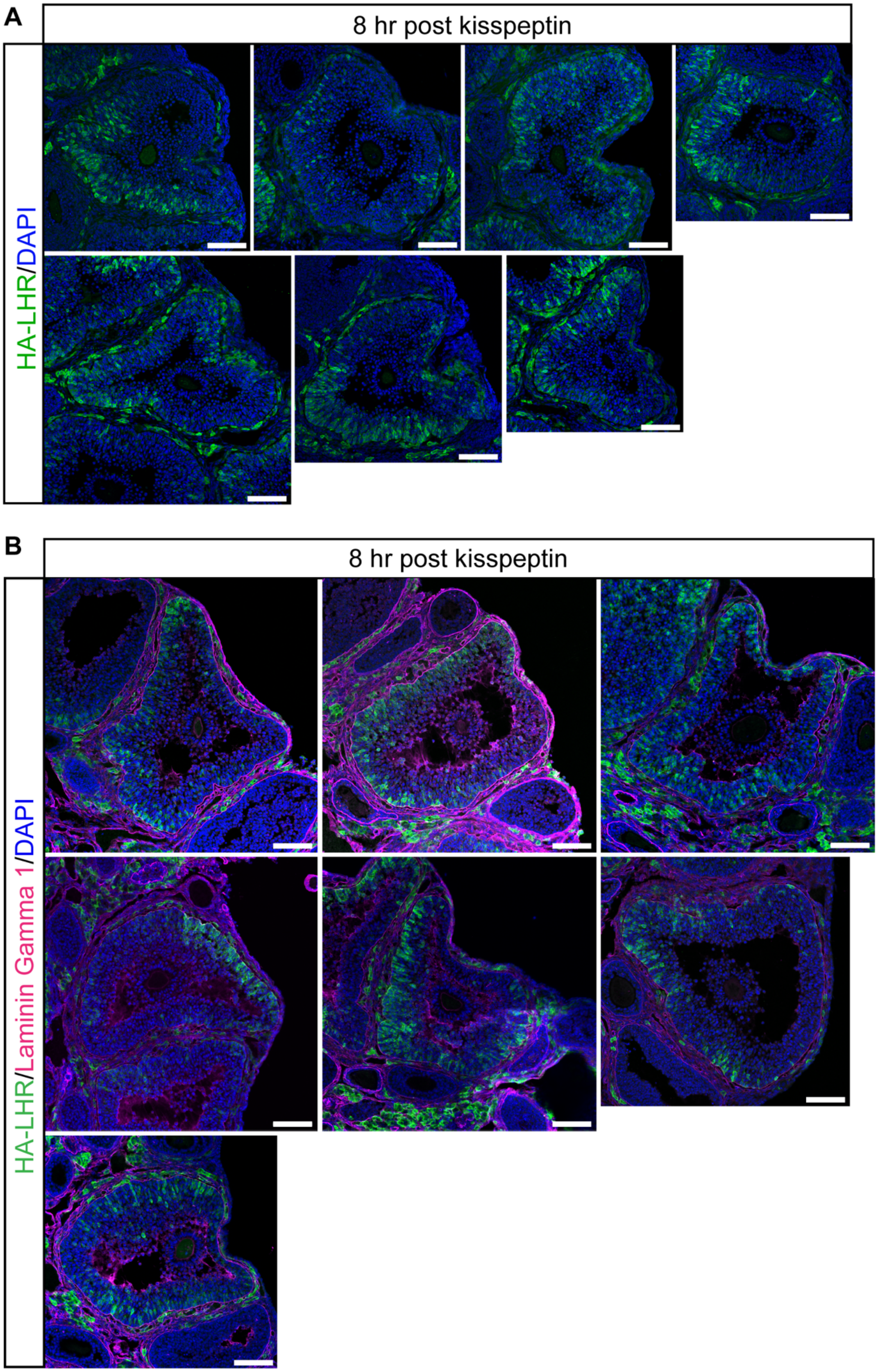

Figure S6

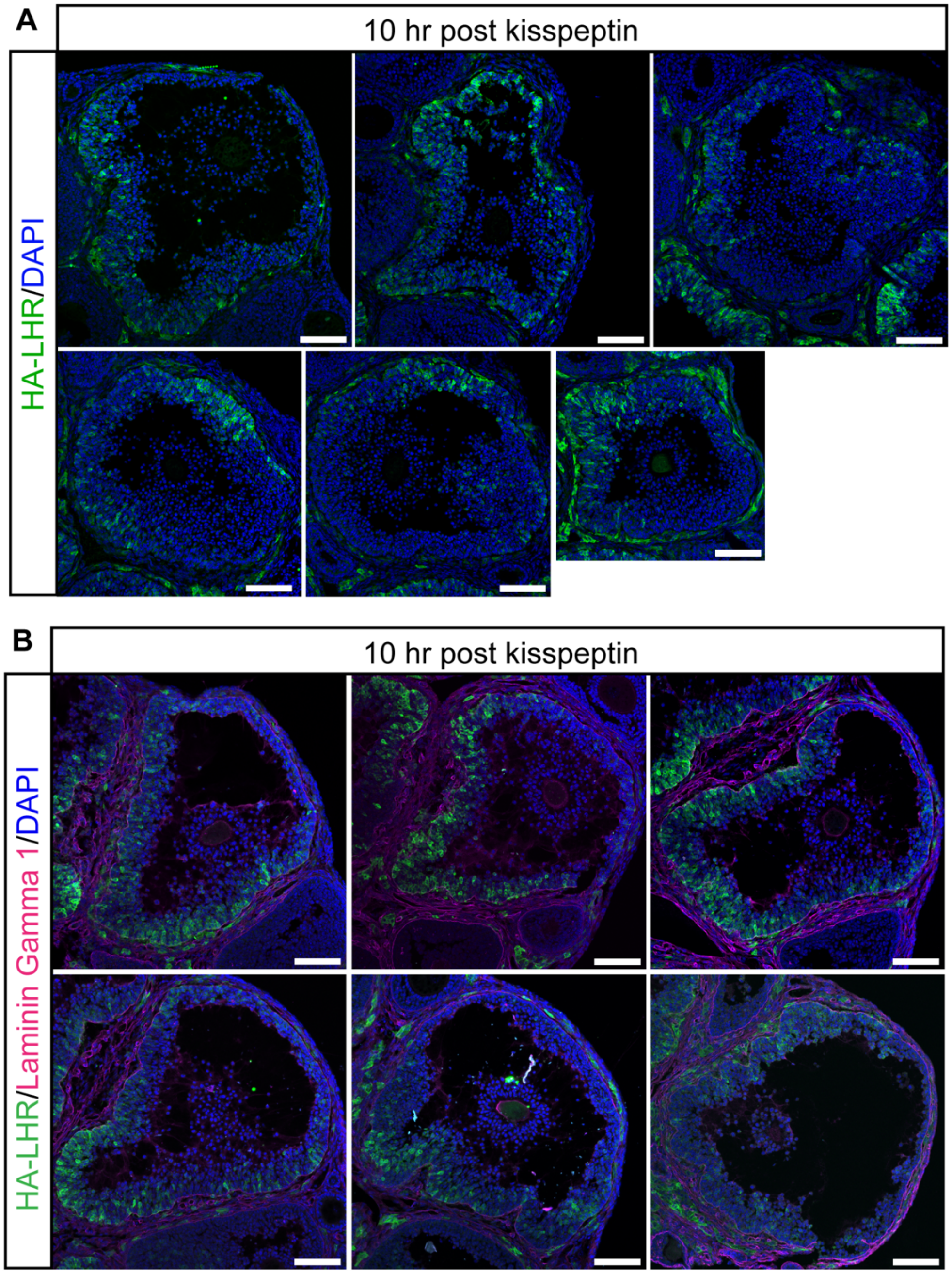
A
10 hr post kisspeptin
HA-LHR/DAPI
B
10 hr post kisspeptin
HA-LHR/Laminin Gamma 1/DAPI

Figure S7

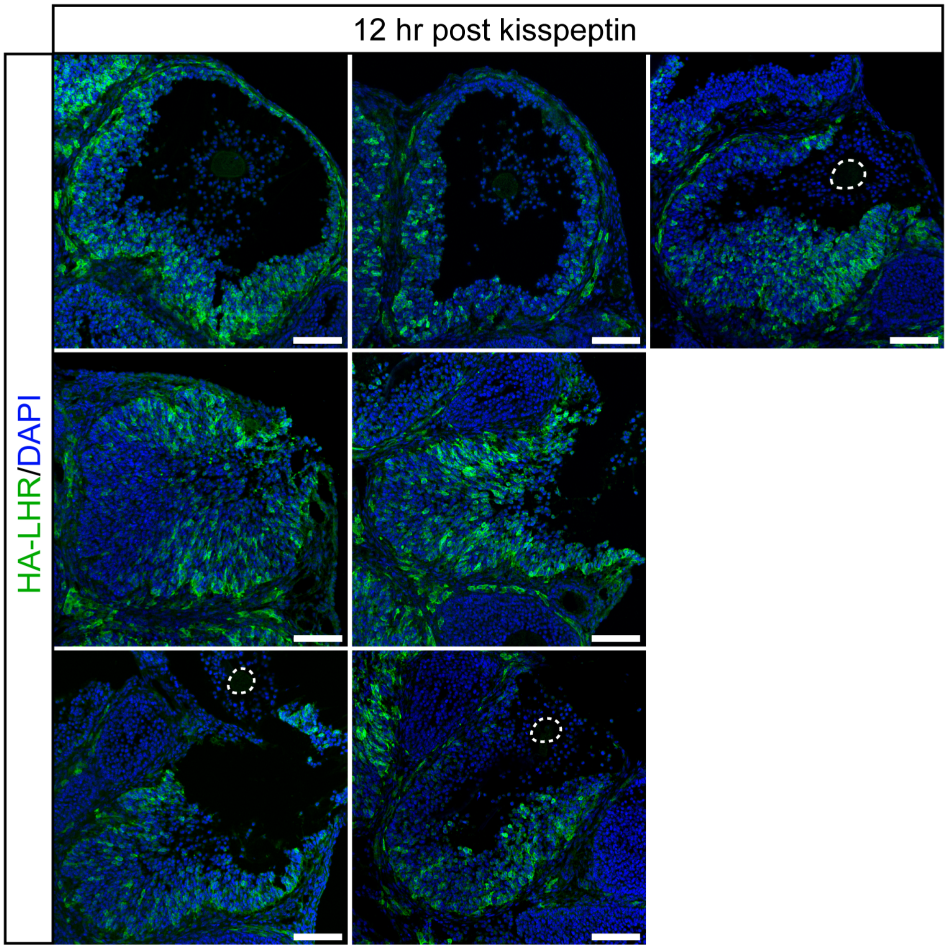

Figure S8

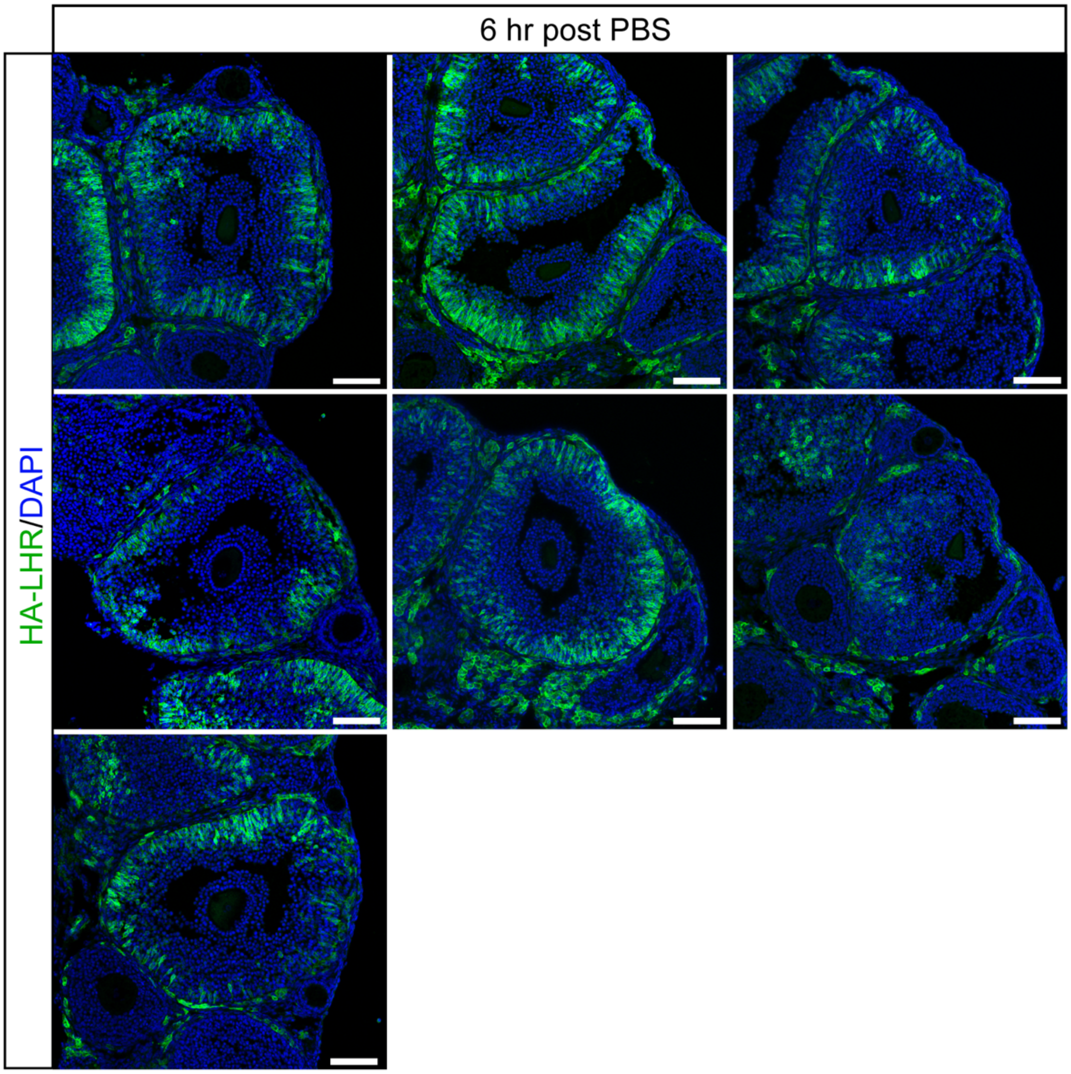

Figure S9

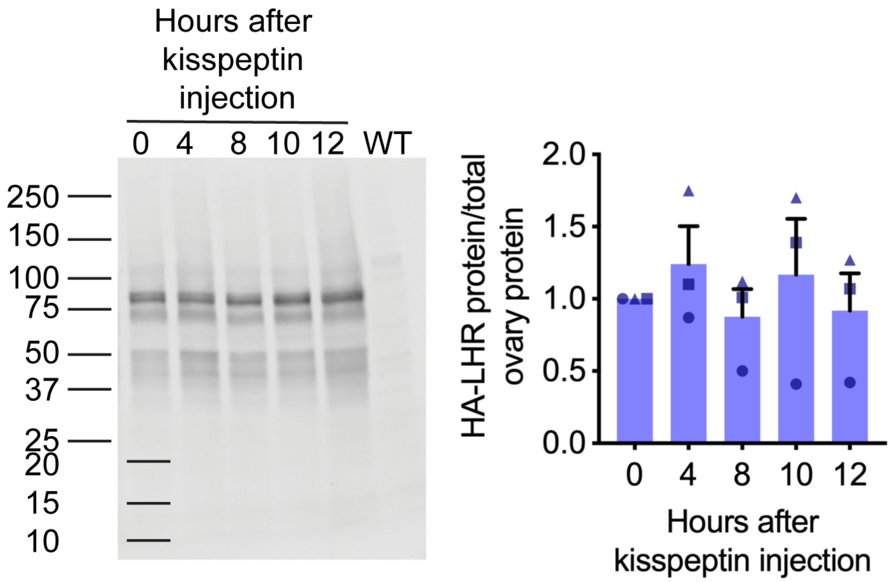
Hours after kisspeptin injection
0
4
8
10
12
WT
250
150
100
75
50
37
25
20
15
10
HA-LHR protein/total ovary protein
2.0
1.5
1.0
0.5
0.0
0
4
8
10
12
Hours after kisspeptin injection

178 Figure S10

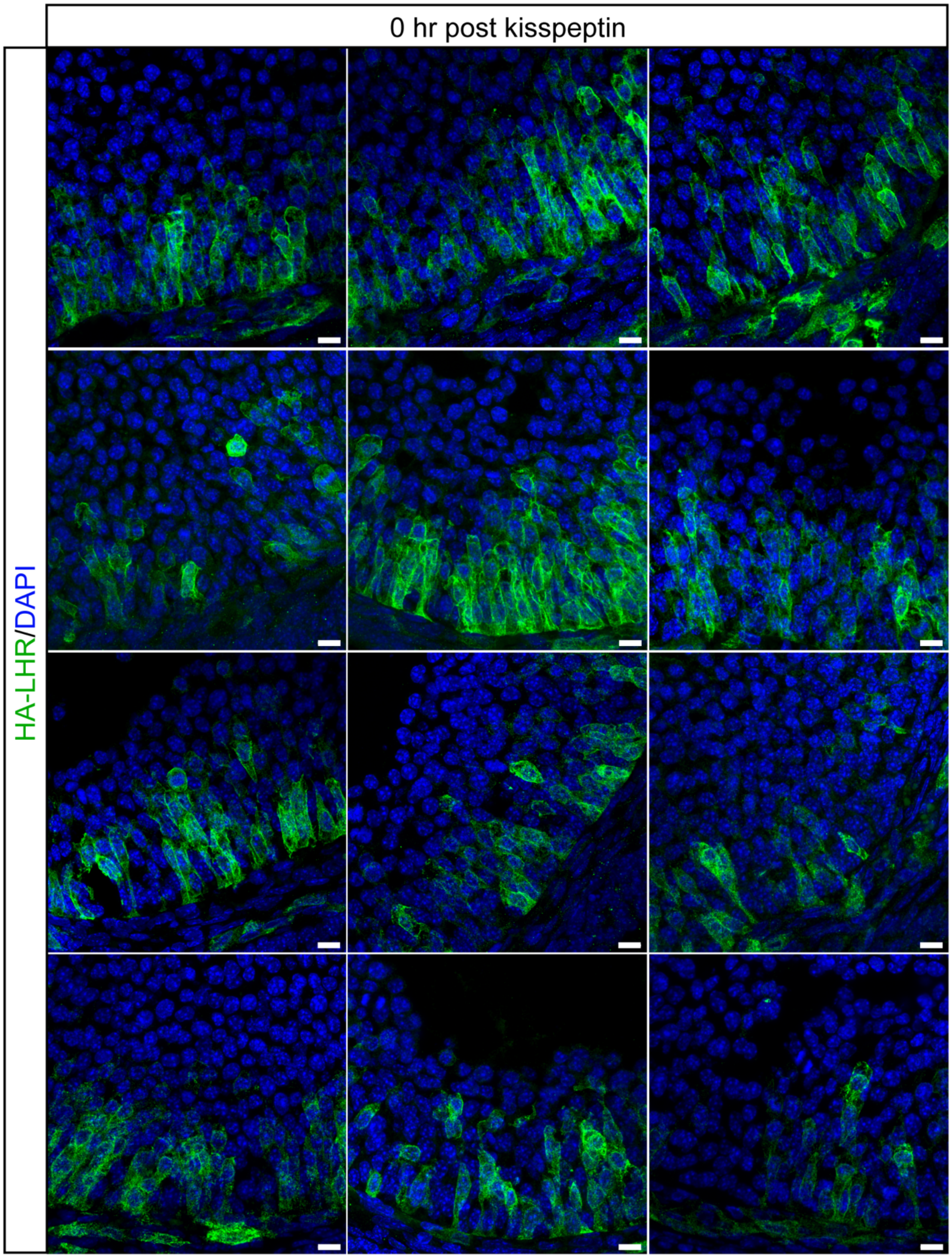


179

180 Figure S11

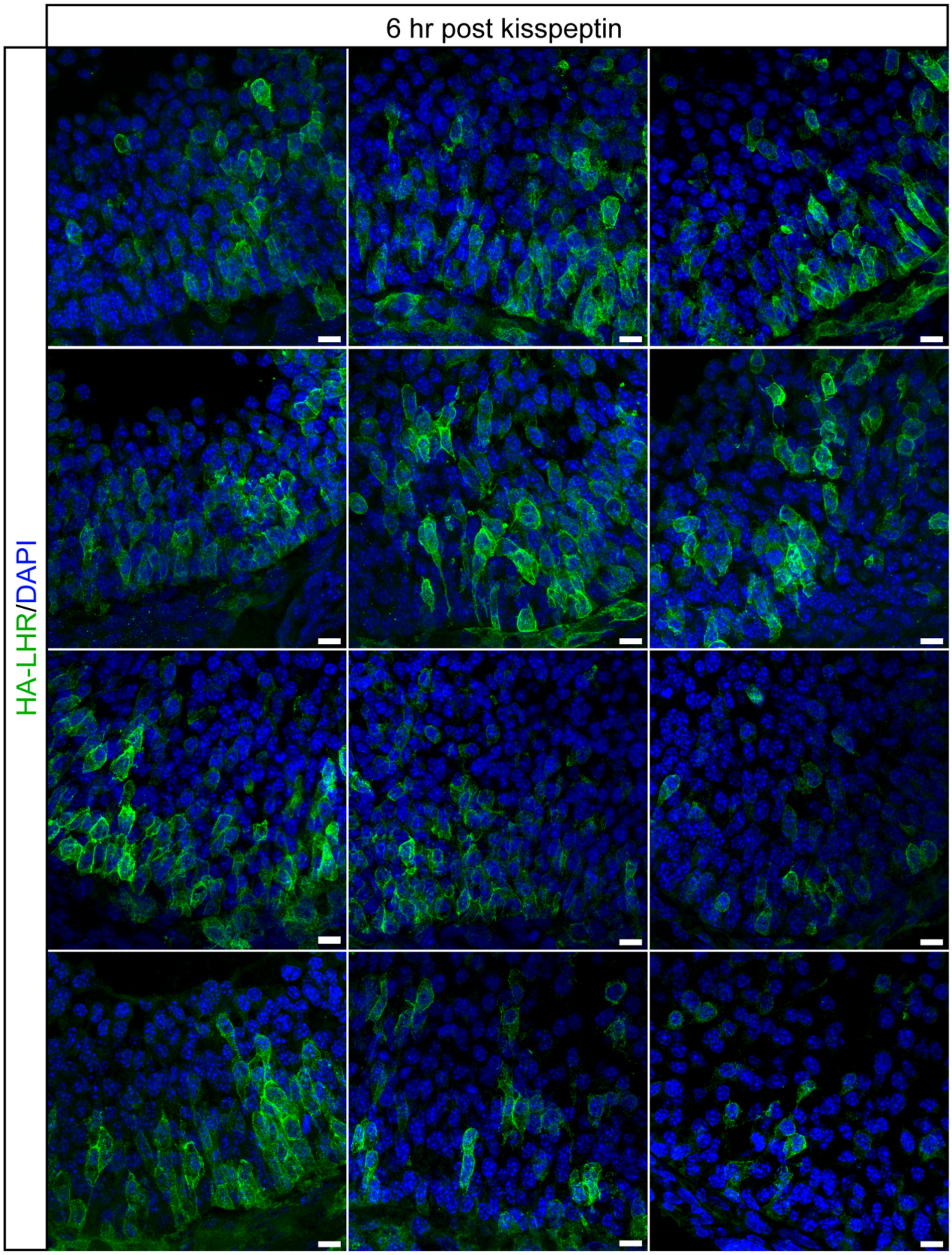


181

182

183 Figure S12

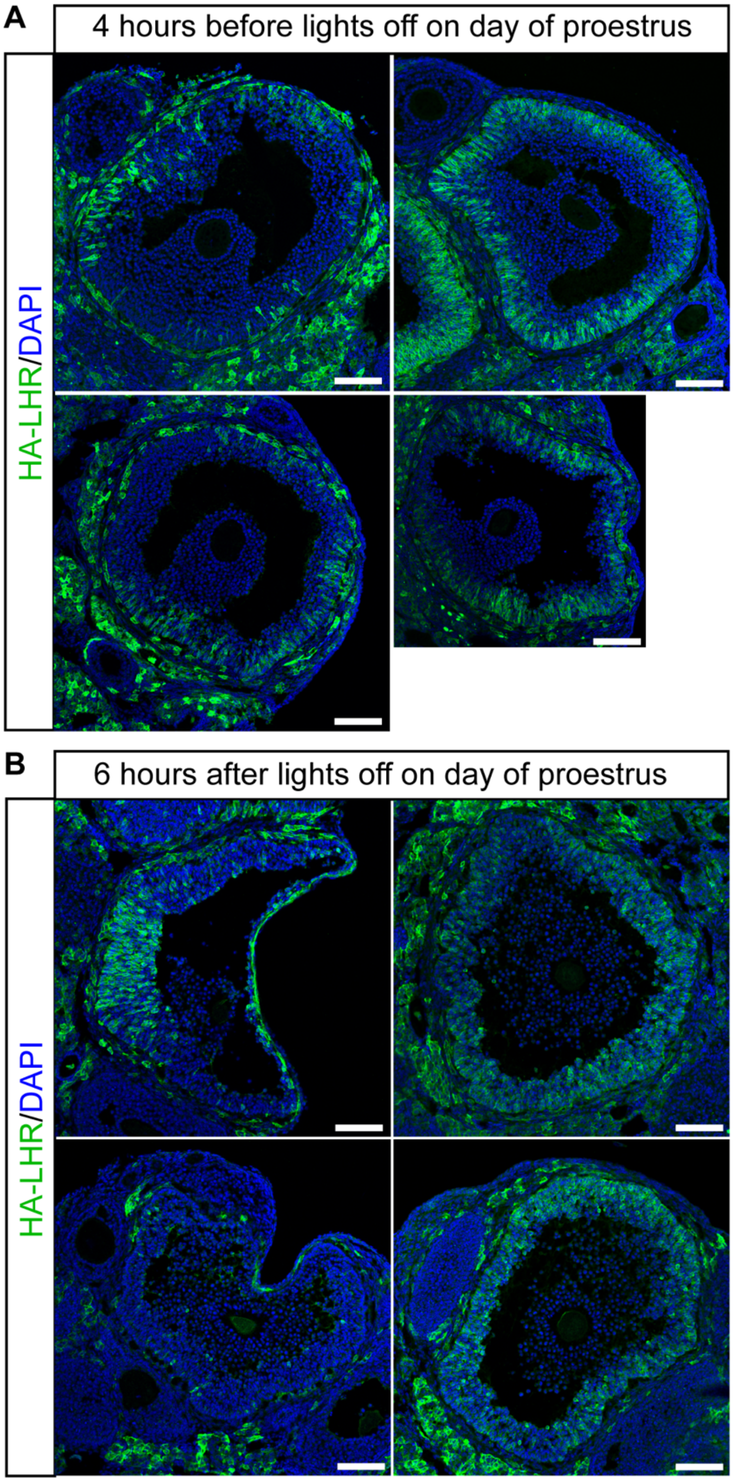


184
